# Molecular stratification of human fetal vaginal epithelium by spatial transcriptome analysis

**DOI:** 10.1101/2024.03.20.585995

**Authors:** Ziying Ye, Peipei Jiang, Qi Zhu, Zhongrui Pei, Yali Hu, Guangfeng Zhao

## Abstract

The human vaginal epithelium is a crucial component in numerous reproductive processes and serves as a vital protective barrier against pathogenic invasion. Despite its significance, a comprehensive exploration of its molecular profiles, including molecule expression and distribution across its multiple layers, remains elusive. In our study, we undertook a spatial transcriptomic analysis within the vaginal wall of human fetuses to fill this knowledge gap. We successfully categorized vaginal epithelium into four distinct zones based on their transcriptomic profiles and anatomical features. This approach unveiled unique transcriptomic signatures within these regions, allowing us to identify differentially expressed genes and uncover novel markers for distinct regions of the vaginal epithelium. Additionally, our findings have highlighted the varied expression of KRT genes across different zone of the vaginal epithelium, with a gradual shift in expression patterns observed from the basal layer to the surface/superficial layer. This suggests a potential differentiation trajectory of human vaginal epithelium, shedding light on the dynamic nature of this tissue. Furthermore, abundant biological processes were found to be enriched in the basal zone by the KEGG pathway analysis, indicating an active state of the basal zone cells. Subsequently, the expression of latent stem cell markers in the basal zone were identified. In summary, our research provides crucial understanding of human vaginal epithelial cells and the complex mechanisms of the vaginal mucosa, with potential applications in vaginal reconstruction and drug delivery, making this atlas a valuable tool for future research in women’s health and reproductive medicine.

## Introduction

In women of reproductive age, the vaginal epithelium is a unique, estrogen-sensitive, non-keratinized stratified squamous epithelium that rests atop the lamina propria. This epithelium is composed of four distinct layers: a cuboidal basal layer, a mitotically active parabasal layer, a glycogen-rich intermediate layer, and a flattened non-cornified superficial layer characterized by pyknotic nuclei [1]. This complex structure serves a pivotal function within the female reproductive system, acting as a crucial barrier against pathogen invasions and playing essential roles in reproductive processes such as childbirth, menstruation, and intercourse [2]. Remarkably, the vaginal epithelium has also been reported to exhibit high permeability [1, 3], a characteristic that has been harnessed for the delivery of medications. This unique property makes the vaginal epithelium a promising target for drug delivery systems, as demonstrated in recent studies [4, 5].

Despite its pivotal role in female reproductive health, our comprehension of the human vaginal epithelium remains incomplete. Previous investigations into its characteristics primarily relied on histological and immunohistochemical (IHC) techniques [1, 6-8], which identified key markers but not provide sufficient dynamic spatiotemporal information of the epithelium’s differentiation process. In recent years, genetic rodent models leveraging advanced gene editing techniques, alongside surgically induced ovariectomy (OVX) models in mice and rats, have shed new light on the differentiation mechanisms of the female reproductive tract epithelium [9-11]. These studies have been complemented by technological advancements such as single-cell RNA sequencing (RNA-seq) and lineage tracing, which have revealed that NGFR^+^Axin2^+^ cells residing in the basal layer of the mouse vaginal epithelium undergo expansion to eventually form the entire epithelium[12]. The maintenance of vaginal epithelial homeostasis is a delicate balance achieved through continuous stem cell development, culminating in shedding or degradation of cells by the immune system [13, 14]. While these rodent models provide valuable insights, it is essential to acknowledge the inherent differences between rodent and human vaginal epithelia [15]. Therefore, a comprehensive and systematic study of the human vaginal epithelium is imperative to elucidate whether a similar developmental pattern exists, specifically regarding the continuous development of vaginal epithelial cells from the basal to the superficial layer.

With the advent of sophisticated sequencing techniques, researchers have delved into the intricacies of the vaginal wall, utilizing transcriptome sequencing to compare patients suffering from vaginal atrophy with healthy women. Their investigations have shed light on the influence of estradiol (E2) on the vagina and its associated gene regulatory networks [16, 17]. Nevertheless, a caveat of this bulk RNA sequencing approach is its inability to provide a nuanced portrayal of epithelial cells. Prior efforts employing single-cell transcriptome sequencing of the vaginal wall in women with pelvic organ prolapse (POP) primarily centered on unveiling pathological mechanisms, leaving much to be desired regarding epithelial characterization[18, 19]. In essence, while prior research has offered incremental insights into this multifaceted organ, a comprehensive understanding of vaginal epithelial cells across various locations and differentiation stages remains elusive.

Keratin (KRT), a prominent cytoskeletal protein found in epithelial cells, serves not only as a structural component but also plays a significant role in cell proliferation, differentiation, and the maintenance of vaginal epithelial polarity and homeostasis [20]. Furthermore, as a type of epithelium, the vaginal epithelium has been reported to abundantly express keratins. The expression of keratins is known be differentiation-dependent, tissue-specific, and often occurs in pairs [21]. Specifically, keratins 6, 7, 8, 10, 14, and 19 have been reported to exhibit distinct temporal and spatial dynamics during the development of the human female reproductive tract [6]. The presence of specific keratins can serve as markers indicating the differentiation status and maturity levels of various epithelial cells [20].

The advent of spatial transcriptomics has ushered in a new era of expression profiling, enabling the interrogation of highly specific regions within tissue sections, even down to subcellular levels. This technology facilitates the correlation of expression data with morphological features, tissue type, and spatial relationships to other structures, thereby offering a more nuanced understanding of multiple tissues and organs[22]. Since there are no spatial transcriptomic data for the vagina, and to construct an unbiased spatial transcriptomic atlas of human vaginal epithelium, our study focused on collecting tissue samples from two human fetuses, aborted at 22^+5^ and 22^+6^ post-conceptual weeks (PCW). This choice was informed by evidence suggesting that the vaginal stratified squamous epithelia of 20-week-old fetuses have attained maturity morphologically [7, 23]. However, it is unknown whether the fetal vaginal epithelium is mature at the molecular level. Both samples were carefully selected as cross-sections of the fetal vagina proximal to the vaginal orifice. Our findings unveil the intricate molecular traits and functionalities of vaginal epithelial stratification, emphasizing the profound cellular diversity that exists within this tissue. This groundbreaking discovery paves the way for potential advancements in vaginal reconstruction [24-29] and the optimization of vaginal drug delivery systems [4, 5, 13]. As research in this area continues to evolve, the implications of these findings are poised to transform our understanding and treatment of vaginal health conditions.

## Materials and methods

### Tissue processing

Vaginal tissue samples were obtained from human fetal specimens in the second-trimester, following an ethically approved process of selective termination of pregnancy by the Ethics Committee of Nanjing Drum Tower Hospital (Ethics number: 2021-131-02). The estimated fetal age was determined based on the mother’s last menstrual period. Six fetal specimens, derived from human fetuses at 22 to 24 weeks, were meticulously collected, with two specimens designated for spatial transcriptomics sequencing, one specimen for determining the time of permeation and three specimens allocated for staining validation.

The procurement of fetal specimens adhered strictly to ethical standards and was carried out with utmost care and precision. This included the careful separation of the skin of the lower abdomen, the meticulous disconnection of ligaments and attachments of the fetal genitourinary system from the abdominal cavity, and the subsequent access to the fetal pelvic-abdominal cavity. The fetal vaginal wall was carefully dissected from surrounding structures, including the bladder, urethra, and anorectum to ensure the integrity and quality of the tissue samples obtained (Figure S1A).

### Spatial transcriptomics

The tissues harvested from the fetal vaginal region underwent a preservation process that involved freezing in isopentane, followed by embedding in OCT and cryo-sectioning, tailored specifically for use with Visium Spatial slides from 10× Genomics. Afterward, H&E staining was conducted, and brightfield microscope images of the stained tissues were captured, laying the foundation for further investigation. In preparation for library construction, an optimal permeabilization time of 24 minutes was established using the Visium Spatial Tissue Optimization Slide kit (Figure S1B). During this carefully timed permeabilization, RNA molecules were captured by barcoded spots on the Visium slides. Following capture, a high-quality library was constructed and its integrity was verified using the DNA 1000 assay Kit from Agilent Technologies. Quantification, performed via the ABI StepOnePlus Real-Time PCR System (Life Technologies), preceded the initiation of the sequencing process. The cDNA library was then sequenced on the Illumina novaseq 6000 by Gene Denovo Biotechnology Co., Ltd (Guangzhou, China).

### Data processing

Data preprocessing was carried out using Seurat V4.1.0 and Space Range 1.2.1 (10x Genomics) [30]. High-resolution H&E staining histology pictures and raw sequencing reads were fed into the Spaceranger Count pipeline to create feature barcode matrices that captured the spatiality of the data, which was used for downstream analysis. During this process, reads were spliced-aware aligned to the human reference genome (GRCh38), and tissues and fiducial spots were automatically detected and aligned. To minimize interference from the urethra, spots around regions exhibiting elevated expression of Uroplakin 3A (UPK3A) were excluded.

Following normalization with SCTransform, runPCA (nPCs = 50) was applied to the V22W-5 and V22W-6 datasets for principal component analysis (PCA). The datasets were then merged using anchor-based canonical correlation analysis (CCA) integration, with the normalization method “SCT” employed to mitigate batch effects. Subsequently, dimensionality reduction, clustering, and visualization were completed using the functions FindNeighbors (dims = 1:50), FindClusters (resolution = 0.6), and RunUMAP (dims = 1:50).

Differential expression analysis was conducted to identify spatially variable genes within both all clusters and specifically within the epithelium, utilizing the FindMarkers function in the R package Seurat. The ‘test.use = wilcox’ parameter was applied, employing a Wilcoxon Rank Sum test to identify differentially expressed genes between two groups of cells. In our vaginal spatial transcriptomics analysis, the expression value of each gene in a given region or cluster was compared against the rest of the spots using the Wilcoxon Rank Sum test. The selection criteria for Differentially Expressed Genes (DEGs) included an adjust p-value < 0.05 and a log2 fold-change > 1.0 (representing at least a 2-fold difference), guiding subsequent Gene Ontology (GO) analysis [31, 32]. For a more focused exploration within the epithelium, we identified DEGs between the four epithelium zones. Genes meeting the criteria of a adjust p-value < 0.05 and a log2 fold-change > 0.585 (equivalent to at least a 1.5-fold difference) were then used to investigate KEGG pathways (https://www.kegg.jp). All enriched pathways within each epithelial zone were visualized through the creation of bubble plots to enhance interpretation.

### Histology and immunostaining

Fresh fetal vaginal tissue samples were fixed in 4% paraformaldehyde for 24-48 hours at 4℃, followed by paraffin embedding after dehydration and hyalinization. The paraffin-embedded tissues were cut into 5-μm-thick slices and stained with Masson’s trichrome according to the kit instructions (BP028, BASMEDTSCI, China).

In our immunohistochemical (IHC) staining protocol, tissue slices were dewaxed, rehydrated, and subjected to the elimination of endogenous peroxidase with 3% hydrogen peroxide. Subsequently, they underwent heat-mediated antigen retrieval and peroxidase blocking using 2% bovine serum albumin (BSA; Biofroxx, Germany) for 30 minutes at room temperature. After blocking with 2% BSA, they were incubated with primary antibodies overnight at 4°C, followed by HRP-conjugated secondary antibodies (Typng, China; MXB, China) at room temperature for 8 minutes. Antigen signals were visualized using 3’3-diaminobenzidine (DAB; Typng, China), and sections were counterstained with hematoxylin before sealing and microscopic examination (DMi8; Leica, Germany).

For immunofluorescence (IF) staining, the steps preceding IF primary antibody incubation are identical to those preceding immunohistochemical antibody incubation. Tissue slices were incubated with primary antibodies overnight at 4°C and then with secondary antibodies at room temperature for 1 hour in the dark. Nuclei were labeled with DAPI (Abcam, USA). Fluorescent images were captured using a fluorescence microscope (DMi8; Leica, Germany).

Both protocols employed a variety of antibodies from different suppliers, as detailed in Table S1.

To assist in validating the spatial characteristics of certain molecules, we adapted immunohistochemistry (IHC) staining images from the Human Protein Atlas (proteinatlas.org) [33]. These images are available under the Creative Commons Attribution-ShareAlike 3.0 International License, allowing for free adaptation, including remix, transform, and build upon the material for any purpose, even commercially. These including IHC staining of SPINK7 (Human Protein Atlas, https://www.proteinatlas.org/ENSG00000145879-SPINK7/tissue/vagina#img), C15orf48 (Human Protein Atlas, https://www.proteinatlas.org/ENSG00000166920-C15orf48/tissue/vagina#img), PI3 (Human Protein Atlas, https://www.proteinatlas.org/ENSG00000124102-PI3/tissue/Vagina#img), IGSF9 (Human Protein Atlas, https://www.proteinatlas.org/ENSG00000085552-IGSF9/tissue/vagina#img) and TP73 (Human Protein Atlas, https://www.proteinatlas.org/ENSG00000078900-TP73/tissue/vagina#img).

## Results

### Spatial RNA sequencing of the human fetal vagina

To explore the molecular features of the human vaginal epithelium, we employed 10×Genomics Visium Spatial Gene Expression assay. This involved the analysis of tissues derived from two human fetuses (22^+5^ weeks and 22^+6^ weeks), designated as V22W-5 and V22W-6 respectively (Figure 1A). Both tissue sections, obtained from cross-sections of the fetal vagina near the vaginal orifice, underwent optimal permeabilization (Figure S1B) before library construction and sequencing. Hematoxylin and eosin staining (H&E) revealed distinct boundaries between lamina propria (LP) and layered epithelia, though the cellular lamina propria exhibited poor demarcation from the muscularis propria (MP) (Figure 1B). The observed structure closely mirrors that of the vaginal epithelia of reproductive-aged women, aligning with previous findings indicating well-developed vaginal epithelia in 20-week fetuses[7, 23].

**Figure 1.**
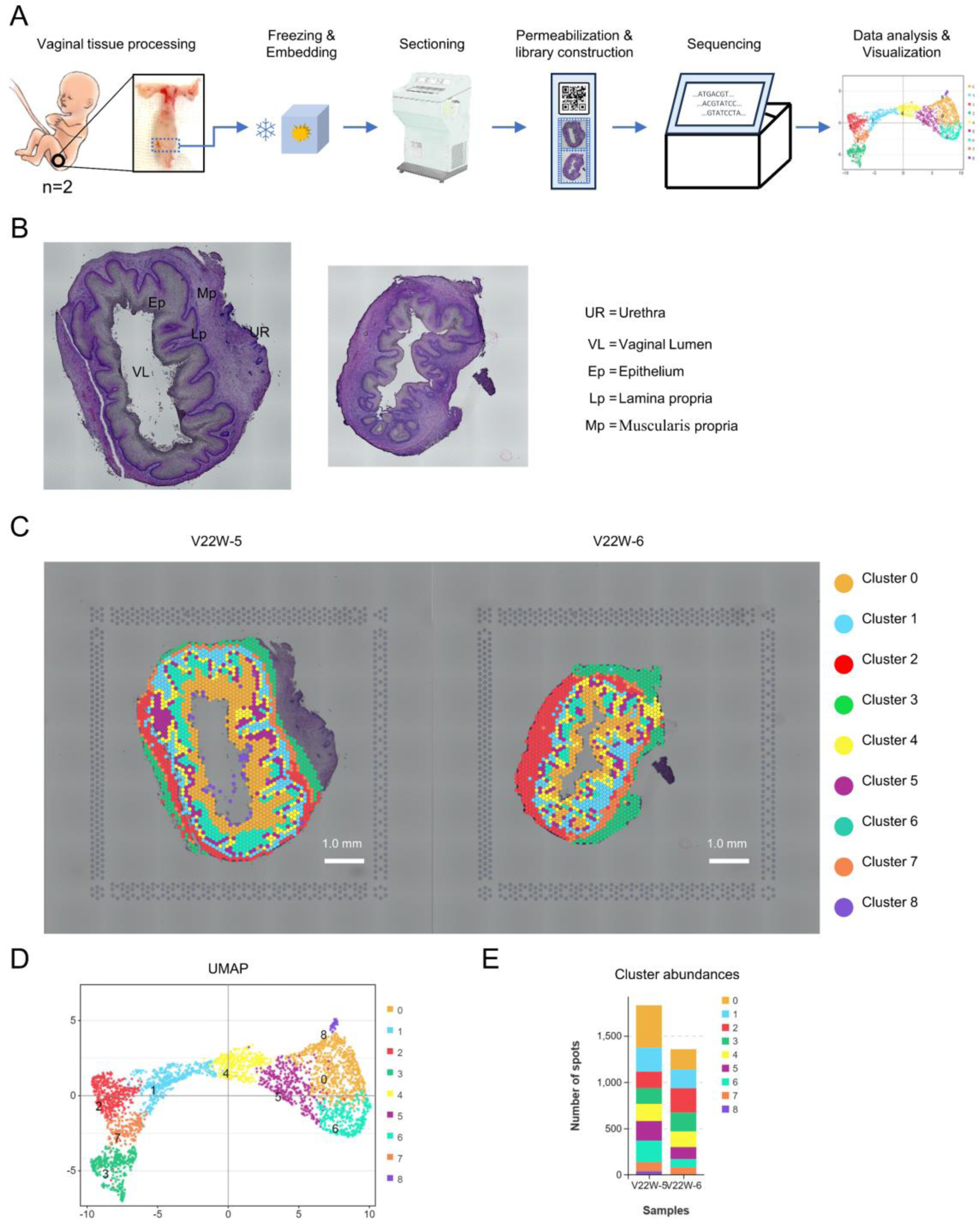
Spatial Transcriptome Analysis of the Human Fetal Vagina. (A) Spatial transcriptomic analysis workflow of the human fetal inferior/lower vagina (n=2, 22W). (B) HE stained cross-sections of the 22PCW fetal vagina near the orifice or introitus. (Left: V22W-5; Right: V22W-6). (C) Spatial transcriptomics results mapped onto the two vaginal sections (Left: V22W-5; Right: V22W-6). (D) UMAP clustering of cell-covered spots from two tissue sections, resulting in 9 unique clusters; spots within the same cluster were color-coded identically. (E) Cluster abundance analysis of V22W-5 and V22W-6 based on spot numbers.

To ensure data integrity, we took precautions to minimize interference from the urethra, which can displace the muscularis propria along the anterior wall of the vagina (Figure 1B, UR). Specifically, we excluded spots exhibits elevated expression of the urothelium marker Uroplakin 3A (UPK3A) [6] and their adjacent regions (Figure 1B, C, and Figure S1C). Additionally, spots located outside the vaginal wall were also excluded from analysis (Figure 1B and 1C). The resulting dataset encompassed 3,187 spots, with 1,831 from V22W-5 and 1,356 from V22W-6. The median number of genes per spot was 4,337 for V22W-5 and 6,032 for V22W-6. Each spot is expected to encompass approximately 1-10 total cells, depending on their composition. Overall, 24,253 genes were identified in the V22W-5 vaginal section, and 25,400 genes were identified in the V22W-6 vaginal section.

After normalization, integration, and dimensionality reduction, the spots were clustered using a resolution of 0.6 yielded cluster closely aligned with the anatomical features of the vagina. The uniform manifold approximation and projection (UMAP) containing cells from the two vaginal sections, revealing 9 distinct clusters (Figures 1D and 1E), and illustrating their relative spatial relationships. Spots within the same cluster, sharing similar transcriptomes, were labeled with the same color, maintaining consistency across both vaginal sections (Figure 1E). Based on morphological and spatial location within the slices (Figure 1C), we assigned the 9 clusters to specific regions: epithelia (clusters 4, 5, 6, 0, and 8), lamina propria (cluster 1), and muscularis propria (clusters 2, 7) and cluster 3. We designated these clusters as the shed vaginal epithelial cells (cluster 8), superficial epithelium zone (Superficial Z, cluster 0), intermediate epithelium zone (Intermediate Z, cluster 6), parabasal epithelium zone (Parabasal Z, cluster 5), basal epithelium zone (Basal Z, cluster 4), the lamina propria (cluster 1), the ventral muscularis propria (Cluster 2) located in the anterior vaginal wall, the dorsal muscularis propria (Cluster 7) situated in the posterior vaginal wall, and Cluster 3. As the histological region of cluster 3 remains unknown and is not the focus of our study, we have refrained from naming it and continue to refer to it as cluster 3. The identification of distinct regions and spatial patterns in the distribution of diverse cell types throughout the vaginal wall, especially within the epithelium, suggests the existence of unique cellular subpopulations with potentially specific functions. Furthermore, the observed asymmetric allocation of clusters, particularly evident in the anterior and posterior vaginal walls, underscores a significant location-dependent heterogeneity within the tissue. This heterogeneity, characterized by variations in cell type composition across clusters, points to distinct microenvironments within the vaginal wall, thereby enhancing our nuanced comprehension of its cellular landscape.

### Spatially variable features and cluster-specific molecular markers revealed by differentially expressed genes in fetal vagina

The identification of distinct regions and spatial distribution of diverse cell types across the vaginal wall suggests unique functions. Spatially variable features in molecular profiles enable the capture of nuanced characteristics within these clusters. To achieve this, we first identified the differential expression genes (DEGs), analyzed the top 10 DEGs (Figure 2A), and then clustered the DEGs according to their unique gene signatures.

**Figure 2.**
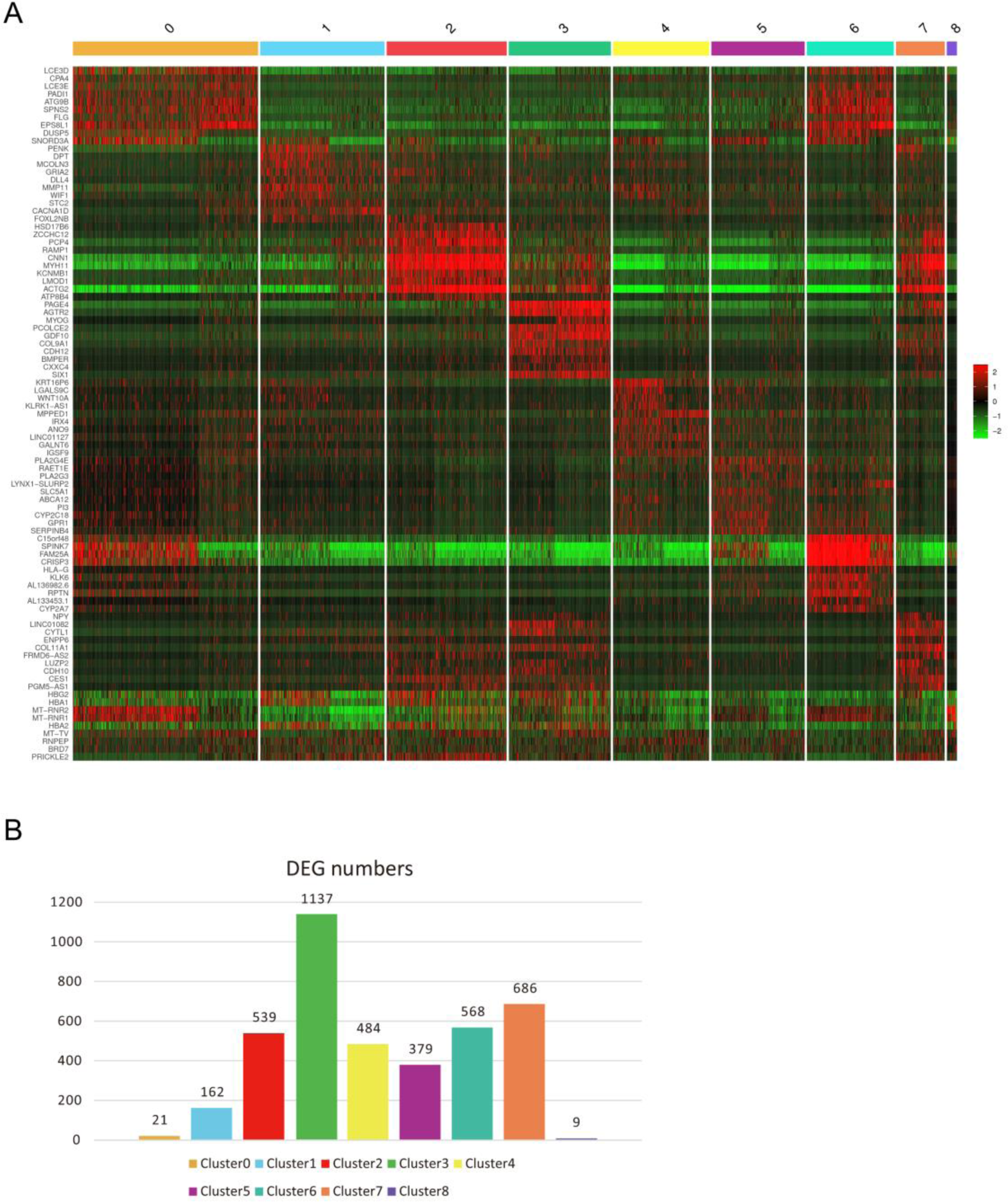
Spatially Variable Features of the Human Fetal Vagina Revealed by Differential Gene Expression Analysis. (A) Heatmap presenting the top 10 Differentially Expressed Genes (DEGs) per cluster, sorted by log2FC. For clusters with fewer than 10 genes, all DEGs are displayed. (B) Variability in the number of DEGs across different cluster.

Among the nine clusters, the superficial zone (cluster 0) exhibited the lowest number of DEGs, with only 21 identified, in comparison to the other epithelial regions, except for the shed cells (Figure 2B). Cluster 0, which encompasses the superficial layers of the vaginal epithelium and serving as the interface between the host and environment. Within this specific zone, Late Cornified Envelope 3D (LCE3D), Late Cornified Envelope 3E (LCE3E), and filaggrin (FLG) exhibited heightened expressions (Figures 2A and 3A), which are indicative of terminally differentiated cells in the squamous epithelium of the epidermis [34, 35]. Moreover, IHC staining of LCE3E and FLG in fetal samples revealed predominant expression in the superficial layers (Figure 3B). This observation supports the notion that the superficial zones of the vaginal epithelium comprise terminally differentiated cells undergoing keratinization in the squamous epithelium, poised for shedding [12, 13]. Moreover, our IHC staining of FLG aligns with previous reports on adult women (https://www.proteinatlas.org/ENSG00000143631-FLG/tissue), suggesting that at 22 weeks, the vaginal epithelium is mature in squamous differentiation and already exhibits similarities to that of adult women. Furthermore, these DEGs in the superficial zone also display high expression levels in the intermediate epithelial zone (cluster 6) (Figure 2A and 3A), implying shared characteristics between these populations and suggesting a gradual reduction in transcript levels from the intermediate to the superficial cells.

**Figure 3.**
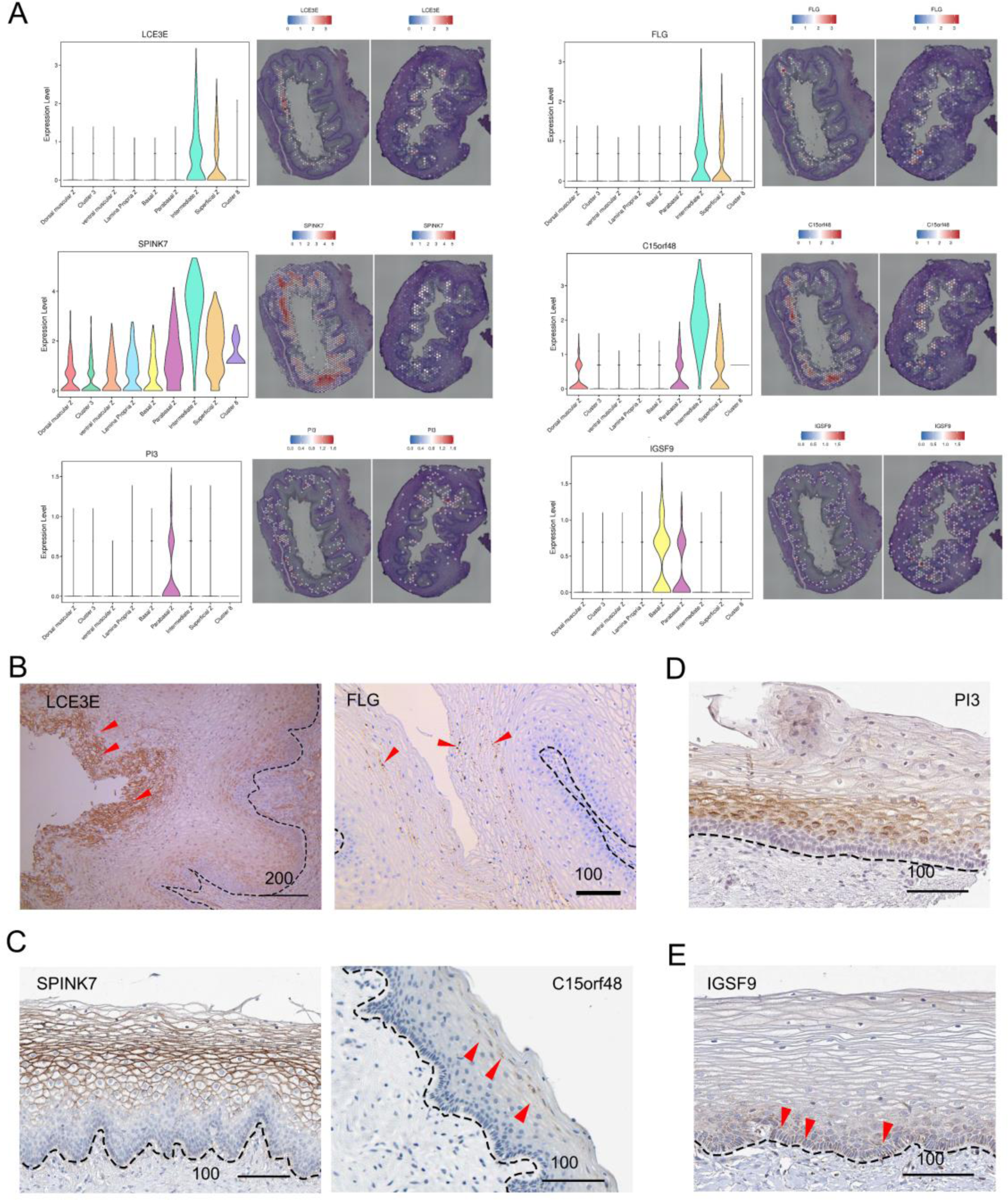
Zone-Specific marker Expression in the Human Fetal Vaginal Epithelium. (A) Left: Violin plot depicting the spatial distribution of DEGs, including LCE3E, FLG, SPINK7, C15orf48, PI3, and IGSF9. Right: Spatial mapping further illustrates the specific zones where these genes are predominantly expressed. (B) Immunohistochemistry (IHC) staining of LCE3E(red triangle arrow) and FLG (red triangle arrow) in fetal vagina. (C) IHC staining of SPINK7 and C15orf48 (red triangle arrow) in adult vagina. (D) IHC staining of PI3 in adult vagina. Notice the PI3-negative cells in the basal layer. (E) IHC staining of IGSF9 in adult vagina. Scale bars are provided in the panels, with units in micrometers (μm). Notably, SPINK7, C15orf48, PI3, and IGSF9 data were cross-validated using the Human Protein Atlas for additional confirmation (see Methods for details). Scale bar: μm.

In the intermediate zone (cluster 6), SPINK7 (serine peptidase inhibitor, kazal type 7) and C15orf48 exhibited significantly elevated expression levels compared to other clusters (Figures 2A and 3A). This spatial expression feature was consistently observed in IHC staining of adult women (Figure 3C), affirming their candidacy as markers for cells in the intermediate zone. SPINK7 is known to regulate the differentiation, barrier function, and inflammatory responses in esophageal squamous epithelium [36], while C15orf48 plays a role in cytoprotection and modulation of immune responses during inflammation [37]. These findings collectively imply that the intermediate zone may play a role in supporting immunization and pathogen defense, contributing to the maintenance of vaginal homeostasis. [16, 35-38].

In the parabasal zone (cluster 5), peptidase inhibitor 3, PI3 could mark these cells well. Our sequencing results also showed that their RNA expression was highest in the parabasal region of the vaginal epithelium (Figure 2A and Figure 3A), which was consistent with the IHC staining results in adult sample (Figure 3D). In the basal zone (cluster 4), immunoglobulin superfamily member 9 (IGSF9) labelled basal zone well (Figure 3E) in adult vagina, as shown in our data (Figure 2A and Figure 3A)

Cluster 1, identified as the lamina propria, provides a unique microenvironment crucial for vaginal epithelia proliferation, differentiation and maintenance [9, 12, 38-42]. It is distinguished from the vaginal epithelium by a delicate basement membrane primarily composed of extracellular matrix (ECM)[8]. The predominantly upregulated genes in this zone are ECM-associated, including metallophosphoesterase domain containing 1 (MPPED1), matrix metallopeptidase 11 (MMP11), stanniocalcin 2 (STC2), and Delta Like Canonical Notch Ligand 4 (DLL4) (Figure 2A and Figure S2A). Masson staining revealed abundant blue-stained collagen fibers in the subepithelial region, indicating a dense extracellular matrix composition (Figure S2B). The extracellular matrix (ECM) serves as a resilient scaffold for vaginal cells, promoting adhesion, proliferation, and migration [39]. Gene Ontology (GO) analysis of differentially expressed genes (DEGs) indicated a notable angiogenic tendency (Figure S2C). Immunohistochemical staining confirmed angiogenesis, showing CD31, CD34, and von Willebrand factor (VWF) presence [40, 41] (Figure S2D). This aligns with prior studies highlighting the vagina’s rich vascular network complexity [41]. Notably, VWF expression was detected in both basal and parabasal layers of the vaginal epithelium (Figure S2D).

Clusters 2, 3, and 7 exhibit a shared upregulation of genes related to muscle cells, including smooth muscle marker actin gamma 2 (ACTG2), calponin 1 (CNN1), myosin heavy chain 11 (MYH11), and Purkinje cell protein 4 (PCP4) (Figure 2A and Figure S3A). Additionally, neuropeptide Y (NPY) is uniquely expressed in cluster 7, corresponding to the muscularis propria located in the anterior wall of the vagina (Figure 2A). This discovery has drawn our attention to the nervous system in the vagina, which is responsive to both mechanical and chemical cues, regulating functions such as blood flow, lubrication, tissue integrity, and playing a crucial role in reflexes related to parturition [43]. Our assessment of neural-related marker molecules, such as vasoactive intestinal polypeptide (VIP), the pan-neuronal marker PGP9.5 (UCHL1), nerve growth factor receptor (NGFR), and S100 calcium-binding protein B (S100B), reveals their predominant expression in the muscularis propria (clusters 2 and 7) and cluster 3 (Figure 2A and Figure S3B).

### Cross-validation of cluster assignment through region-specific markers and histological characteristics

Furthermore, we integrated key markers identified in previous studies into our analysis, mapping them onto our spatial transcription data. We organized these markers, indicative of distinct vaginal characteristics, into a heatmap for better visualization (Figure 4A). Additionally, spatial characteristics observed on the H&E section slices were also incorporated as constraints to guide the cluster assignment process (Figure S4A). In the majority of cases, we found that well-established cell type markers reliably identified specific cell types within vaginal tissue. For instance, AQP1 marked the subepithelial area, aligning with capillaries and venules, as reported previously [44]. AQP3 was observed in the epithelial zone, consistent with its known role as an epithelium marker [44]. Similarly, ACTA2[45] and Desmin (DES) [46] in the muscularis propria confirmed their utility in identifying smooth muscle cells. Additional markers, including ISL1 for vaginal mesenchyme [6], LYVE1 for lymphatic vessels [47], Tp63 for basal region cells of the vaginal epithelium [6], and F11R for the parabasal and basal layers of the epithelium [8], were observed in corresponding locations with related patterns (Figure 4A and Figure S4A). Overall, these findings highlight the robustness and precision of these markers in characterizing distinct cellular components within the vaginal tissue, further confirming our assignment of clusters to corresponding zones of the vagina.

**Figure 4.**
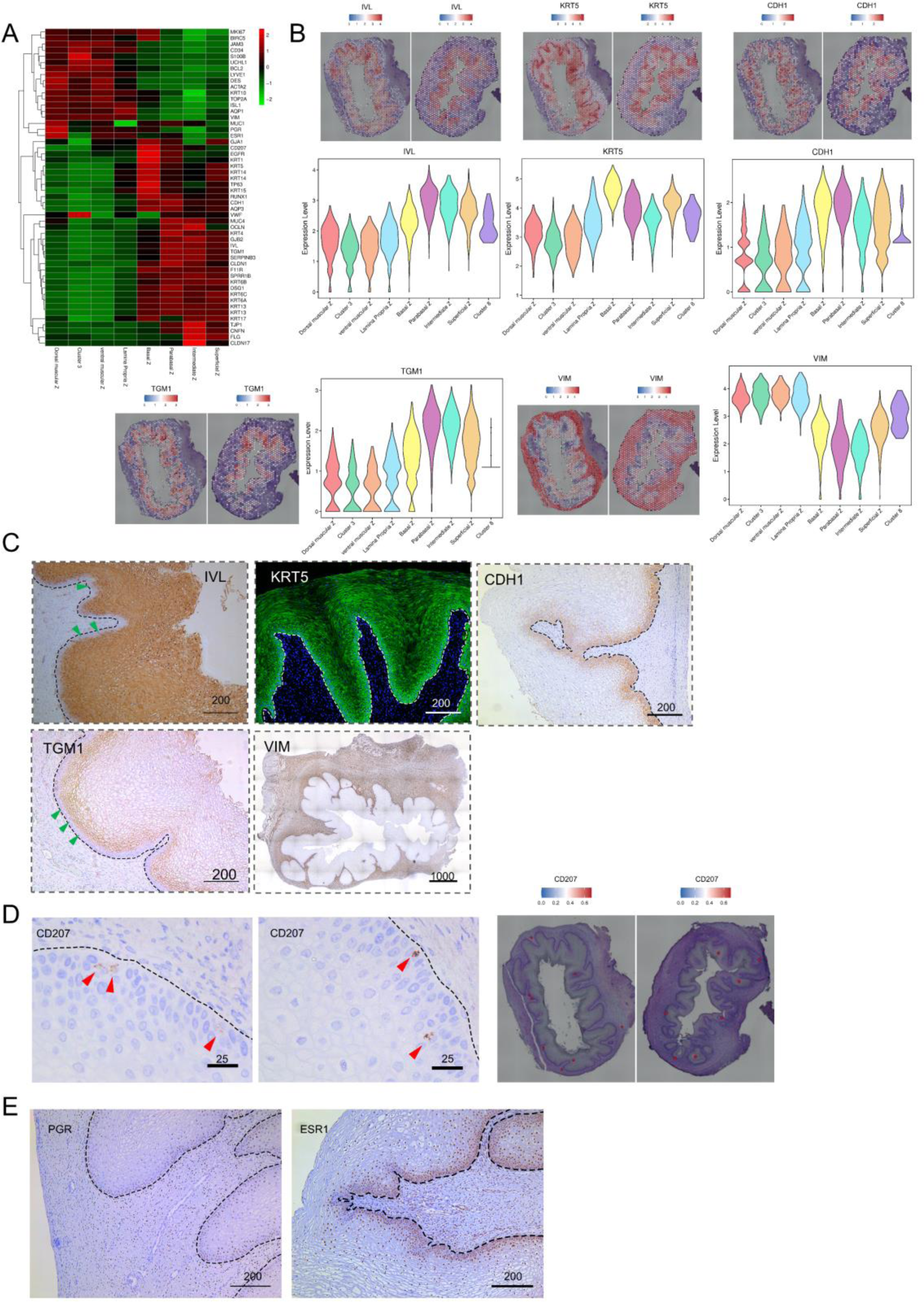
Cluster Assignment using Region-Specific Markers and Constraints from Histological Characteristics. (A) Heatmap displaying the relative expression levels of canonical markers in each cluster. (B) Spatial distribution features of markers (IVL, KRT5, CDH1, TGM1, and VIM) in the human fetal vagina illustrated through spatial mapping and violin plots. (C) Verification of markers expressed in vaginal epithelia through immunofluorescence (KRT5) and immunohistochemical staining (IVL, CDH1, TGM1, VIM) in fetal vagina. (D) IHC staining (left, middle) and spatial mapping of CD207 in the basal layer (red triangular arrow) in fetal vagina. (E) IHC staining of PGR and ESR1 in the human fetal vagina. Scale bars are included in the panels, with units in micrometers (μm).

To further validate the applicability of the markers chosen for assigning clusters in the vaginal tissue of 22+ week fetus, we conducted IF and IHC experiments for selected markers. Involucrin (IVL), a protein subunit of keratinocyte cross-linked envelopes, is a distinctive marker for suprabasal differentiation in stratified squamous epithelium [48]. IVL is highly expressed in the parabasal region of vaginal epithelium (Figure 4B, adjusted p-value < 0.0001). And this protein makes its appearance in the cells immediately above the basal layer (Figure 4C). At 20 weeks of gestation, the expression of basal cytokeratin 5 (KRT5) emerges and is crucial for maintaining the integrity of squamous epithelium [49]. In our study, KRT5 expression was highest in the basal region (Figure 4B). IF demonstrated intense expression of KRT5 in the fetal vaginal epithelium at 22^+^ weeks, particularly in the basal zone (Figure 4C). The staining pattern observed in IF aligned with the trend of RNA expression obtained from sequencing data. E-cadherin (CDH1) is the common transcellular component of all epithelial adherens junctions, and it exhibits its brightest expression in the parabasal and basal layers of the vaginal epithelium in adult women [8]. Consistent with these findings, our study showed that the CDH1 gene had the highest expression in both our parabasal and basal regions (Figure 4B). Moreover, protein expression of CDH1 was notably strong in the basal layer and parabasal layers (Figure 4C). Transglutaminase 1(TGM1) regulated by estradiol plays a key role in the process of terminal differentiation of rat vaginal epithelial cells [50]. Our sequencing results also showed that its expression was highest and was the DEG in the parabasal region of the vaginal epithelium, which was consistent with the IHC staining results (Figure 4B and 4C). Additionally, the wide expression of the stromal cell marker vimentin (VIM) [51] is notable in non-epithelial areas, and IHC staining for VIM can effectively distinguish between epithelia and mesenchyme (Figure 4C). Taken together, these results suggest that our sequencing outcomes are reliable, our clustering strategy is reasonable, and the gene expression pattern in the 22-week fetal vaginal epithelium closely resembles that of adults.

Since the vaginal epithelium has an important role as an immune barrier, we also examined the expression of several immune cell marker molecules in the sequencing data. Specifically, we observed the expression of CD68 for macrophage [52], which was primarily expressed in the intermediate zone (Figure S4B). Additionally, CD207, a marker for Langerhans cells [53], was expressed in the basal region (Figure 4D). Furthermore, CD45, which is involved in the initiation of T cell receptor signaling [54] was primarily expressed in the basal zone (Figure S4C). These results suggest that immune cells already present in the vaginal epithelium of 22^+^ week fetuses and the expression of immune cells in the vaginal epithelium is location-specific.

Estrogen-stimulated vaginal epithelium expressed ESR1 widely in vaginal mesenchyme and epithelium (Figure 4E). In our fetal samples, the progesterone receptor (PGR) is uniformly expressed in the epithelium of the basal region and widely expressed in the subepithelial region (Figure 4E), suggesting that steroid hormones broadly influence the development of the vagina at this time.

### Keratin genes reveal the heterogeneity of vaginal epithelial cells

The vaginal epithelium, characterized by a diverse array of molecular markers (Figure 4A), demands a targeted examination of pivotal molecules to elucidate their interactions and developmental trajectories. Leveraging insights from studies on keratin expression in human epithelium, especially its pivotal role in differentiation and regulation [55], we analyzed Keratin proteins (KRTs) within the vaginal epithelium (Figure 5A, Table S2) to shed light on the underlying processes. Through our comprehensive spatial transcriptomics analysis of the vagina, we were able to identify a substantial representation of human genomic keratins within the epithelium. Specifically, we detected 42 out of the 56 reported keratins, accounting for an impressive 75% (Table S2). Among these, the most abundantly expressed keratins in the epithelium included KRT 13, 6A, 5, 14, 4, 6C and 19 (Table S2). To unravel the intricacies of epithelial heterogeneity with respect to proliferation and differentiation, we systematically clustered the detected keratins and visualized them in a heatmap (Figure 5A). This analysis provided valuable insights into the specific keratins associated with each zone of the epithelium.

**Figure 5.**
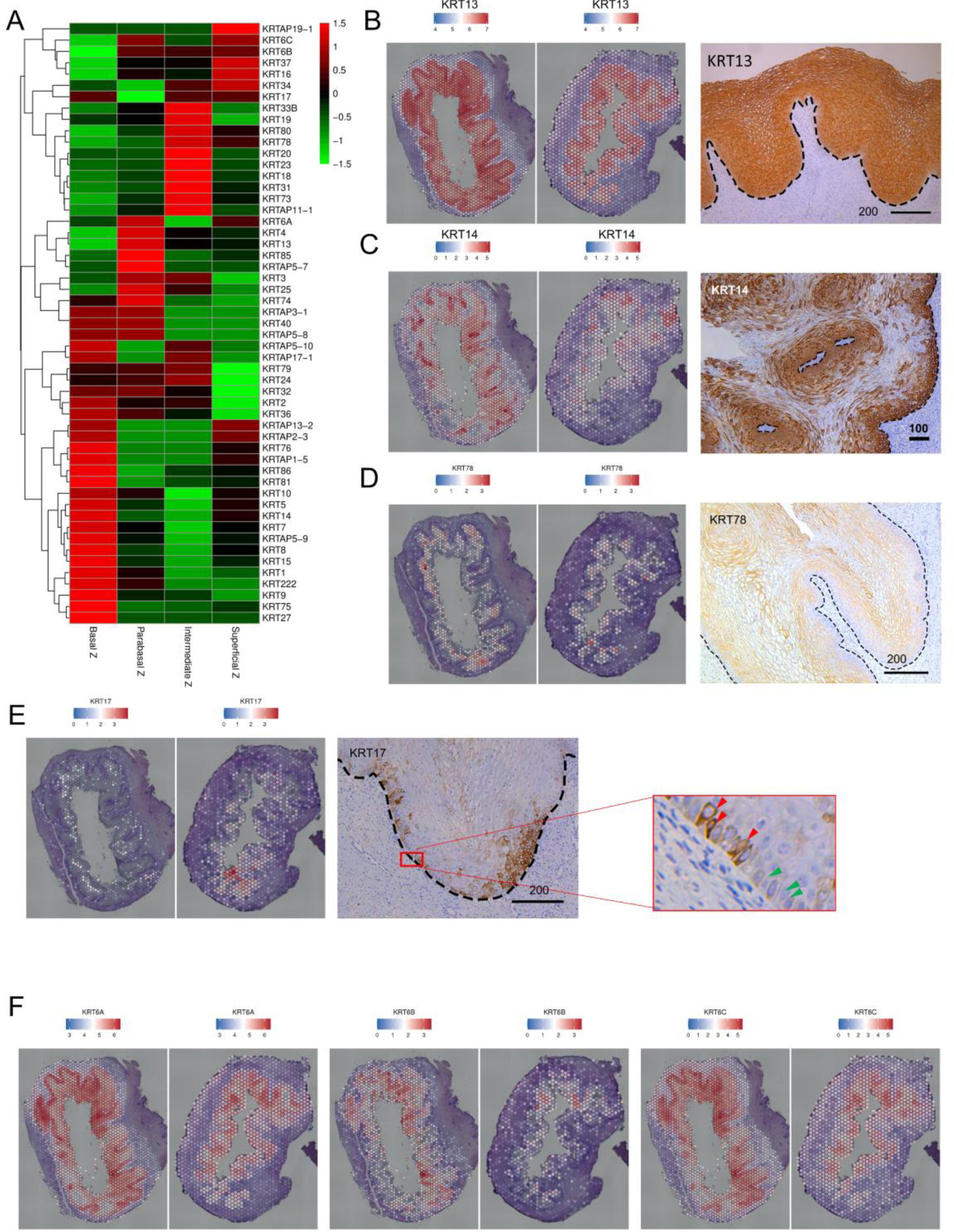
Keratin Genes Analysis Reveals Heterogeneity of Vaginal Epithelium. (A) Heatmap analysis illustrating the zone-specific dominant keratin family genes across the vaginal epithelium. (B) Left: Spatial mapping of KRT13 across the vaginal epithelium. Right: IHC staining of KRT13 in fetal vagina. (C) Spatial mapping and IHC staining of KRT14 in fetal vagina. (D) Spatial mapping of KRT78 and IHC staining of KRT78 in fetal vagina. (E) Spatial mapping of KRT17 and IHC staining of KRT17 in fetal vagina. KRT17 positivity (red arrow); KRT17 negative (green arrow). (F) Spatial mapping of KRT6A, 6B, and 6C. Scale bar: μm.

Our findings revealed distinct expression patterns of keratins across different zones, highlighting spatial heterogeneities. Firstly, we find out that KRT13 showed significantly elevated expression across all zones (Figure 5B), emphasizing its distinct and potentially crucial role in shaping the molecular landscape of the vaginal epithelium. Notably, several KRTs were predominantly expressed in the basal region (Figure 5A). In particular, KRT5/14, which was most abundant in the basal zone, exhibited a gradual decrease in expression towards parabasal and intermediate zones (Figure 4C and Figure 5C), aligning with tissue stratification. Interestingly, a marked increase in KRT5/14 expression was observed in the superficial zone, prompting further exploration of its functional significance (Figure 5A). In contrast to KRT5/14, KRT1/10 demonstrated relatively lower levels across all zones (Table S2). KRT78 was predominantly expressed in the intermediate region and superficial region. This finding confirmed by heatmap analysis, spatial mapping, and further substantiated by IHC, which revealed an ascending pattern from the base to the intermediate region (Figure 5A and 5D). Interestingly, KRT17 expression formed discrete clusters confined to a small region (Figure 5E). In the basal layer, KRT17 was observed in only a subset of cells (Figure 5E, enlarged, red), while others lacked this expression (Figure 5E, enlarged, green), revealing heterogeneity within basal layer. Furthermore, our analysis centered on the three isotypes of KRT6, which, along with KRT16/17, play a crucial role in epithelial healing [56]. These isotypes exhibited wide distribution patterns along the vaginal epithelium (Figure 5F).

Functional enrichment analysis of differentially expressed genes among vaginal epithelial zones

Given our observation of distinct gene expression patterns in various regions of vaginal epithelium, the functions and pathways of these epithelial regions were enriched (Figure 6) by Kyoto Encyclopedia of Genes and Genomes (KEGG). The comprehensive analysis allowed us to delve deeper into the biological roles played by the DEGs (Figure S5A).

**Figure 6.**
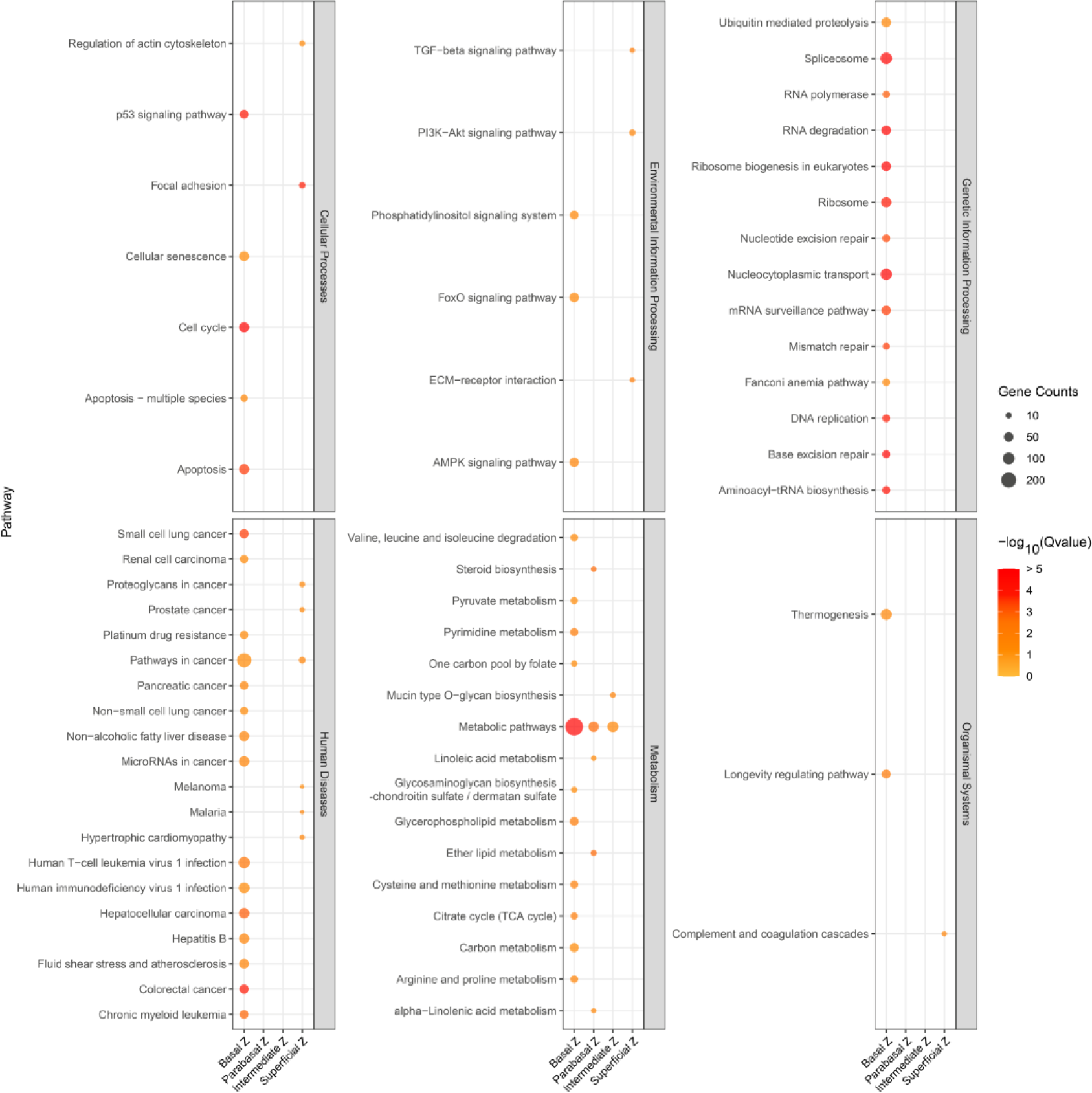
KEGG Analysis Reveals Diverse Pathways in Various Epithelial Zones. Bubble plot presenting all the KEGG pathways to elucidate the pathways active within each region of vaginal epithelia. Colors represent -log10(Q-value), and bubble size indicates the gene count enriched within each pathway.

Notably, the DEGs in basal cell zone were predominantly linked to cell cycle, apoptosis, cellular senescence, p53 signaling pathway, citrate cycle (TCA cycle), and metabolic pathways (Figure 6). Additionally, they were found to be associated with genetic information processing pathways such as Nucleocytoplasmic transport, DNA replication, nucleotide excision repair, mismatch repair, base excision repair, spliceosome, RNA polymerase, ribosome biogenesis in eukaryotes (Figure 6). This suggests that the basal region cells of the vaginal epithelium are highly active and could potentially play a pivotal role in the maintenance, proliferation, and differentiation of epithelial stem cells in this region.

We conducted an examination of several markers identified for the basal zone, along with markers typically associated with stem cells in other organs, to assess their potential as indicators of vaginal epithelial stem cells. In the vaginas of 22+ week fetuses, proliferation was predominantly observed in the 2-3 layers of cells in the basal region, as evidenced by MKI67 staining (Figure 7A). Subsequently, we performed immunohistochemical staining for Axin2 and NGFR, markers of mouse vaginal epithelial stem cells, which are expressed in a single layer of cells in the basal region of the mouse vagina [12, 55]. Consistent with the spatial expression pattern observed in mouse vaginal epithelium, NGFR labeled only one layer of cells in the basal region of the human fetal vaginal epithelium. However, AXIN2 expression was detected in several layers of cells in the basal region (Figure 7A). This pattern differed from that observed in mouse vaginal epithelium, where only a basal layer of positive cells was present.

**Figure 7.**
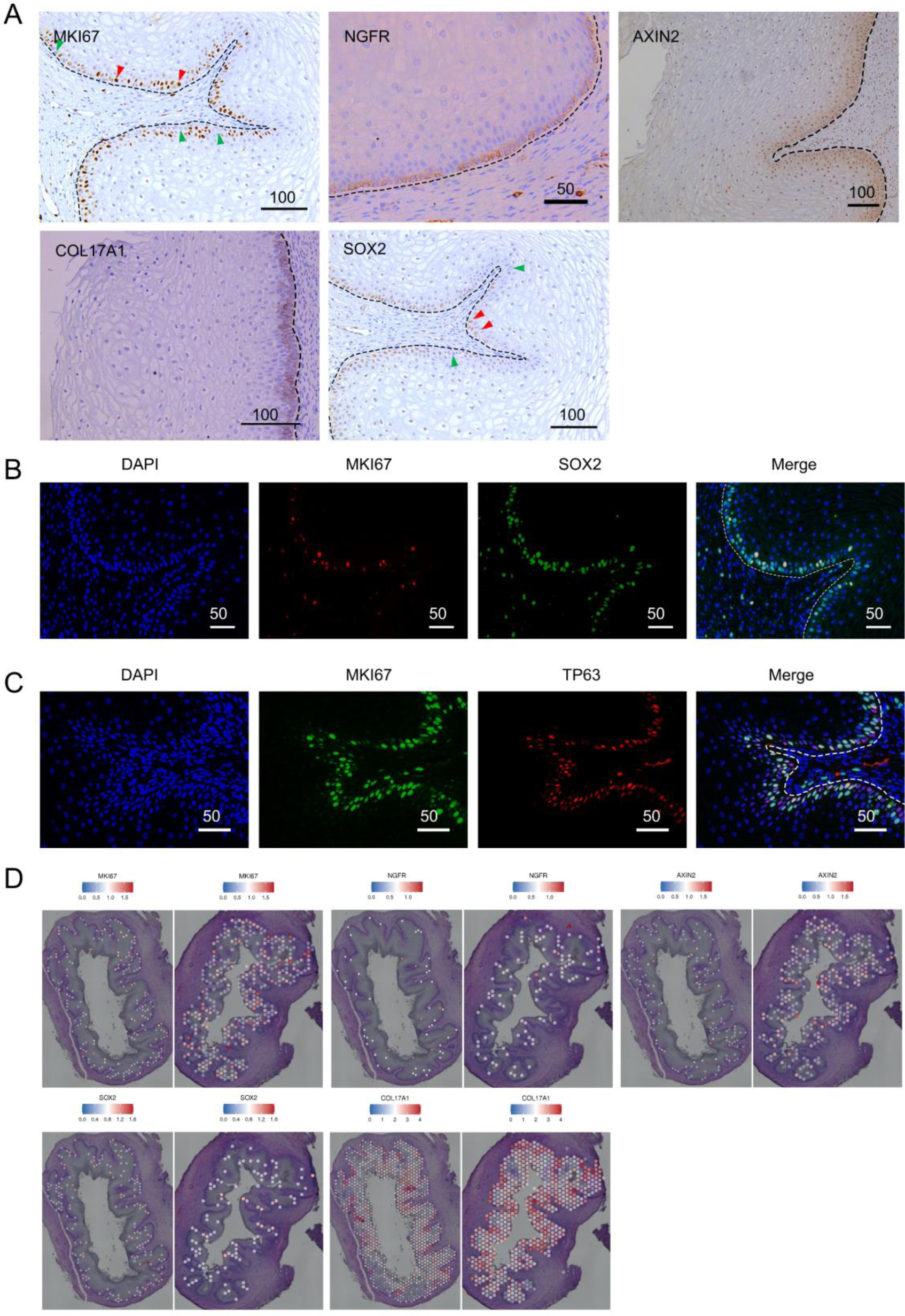
Potential Progenitor Epithelial Cells in the Vagina. (A) IHC staining of proliferation (MKI67) and stem cell-related markers (NGFR, AXIN2, COL17A1, SOX2) in the active basal layer in fetal vaginal epithelium. (B) Immunofluorescence (IF) staining of MKI67 and SOX2 in fetal vagina. (C) IF staining of MKI67 and TP63 in fetal vagina. Scale bar: μm. (D) Spatial mapping of MKI67, NGFR, AXIN2, SOX2 and COL17A1.

Furthermore, we analyzed the expression of known stem/progenitor cell markers from other organs in the fetal vagina. COL17A1, a marker of epidermal stem cells encoding a collagen with a hemibridged granule structure involved in keratinocyte adhesion [56], exhibited differential expression in the basal region of the vaginal epithelium and was specifically expressed in the basal layer (Figure 7A). This expression pattern was consistent with that observed in adults (The Human Protein Atlas. https://www.proteinatlas.org/ENSG00000065618-COL17A1/tissue/vagina). Additionally, we found that SOX2, a key regulator of various stem cell types including embryonic stem cells (ESCs) and neural progenitor cells (NPCs) [57], was predominantly expressed in the basal layer of the epithelium, although not all basal cells were SOX2-positive (Figure 7A). Cells co-labeled with MKI67 and SOX2 immunofluorescence were observed in the basal region (Figure 7B). TP63, a master regulator of stratified epithelia responsible for initiating the stratification program and maintaining the self-renewal capacity of stratified epithelial stem cells [40, 58, 59], was specifically located in the basal region of the vaginal epithelium. Significant overlap was observed between MKI67-positive and TP63-positive epithelial cells (Figure 7C). TP73, a marker of the basal epidermal stem cell populations located near hair follicles in scale-like skin[57], was specifically expressed in the basal region of the vaginal epithelium (Figure S5B and S5C) and was also present in the basal layers of adult vaginal samples (Figure S5D).

Furthermore, the parabasal layer demonstrated a marked increase in substance synthesis and metabolic activity. This zone was found to be abundant in various metabolic pathways, especially about lipids synthesis and metabolism, including Steroid biosynthesis, Linoleic acid metabolism, alpha-Linoleic acid metabolism, Ether lipid metabolism (Figure 6). The intermediate zone showed an enrichment in Mucin type O-glycan biosynthesis (Figure 6). In contrast, the superficial region did not exhibit significant enrichment in metabolic pathways, which could be attributed to the terminal differentiation stage of cells in this region. Collectively, these findings underscore the high metabolic activity within the vaginal epithelium of basal, parabasal and intermediate zones, with epithelial cells metabolizing distinct substances in specific areas.

Importantly, the basal zone also showed an enrichment in diseases related to pathogenic microbial infections, such as human immunodeficiency virus 1 infection, and Human T-cell leukemia virus 1 infection (Figure 6), highlighting the immune function of vaginal epithelial cells. In conclusion, our clustering strategy revealed distinct subgroup of vaginal epithelial cells that are enriched in specific biological processes.

## Discussion

Our study employed the 10×Genomics Visium Spatial Gene Expression assay to conduct a thorough examination of cellular composition and gene expression profiles within the vaginal epithelium of two human fetuses (22+5 weeks and 22+6 weeks). This approach allowed us to unveil the molecular features and heterogeneity across different epithelial layers. By incorporating spatial constraints and region-specific markers, we achieved enhanced accuracy in cluster assignments, revealing nine distinct clusters that aligned with the anatomical features of the vagina, including epithelia, lamina propria, and muscularis propria regions. Furthermore, the epithelium was delineated into four distinct regions: superficial, intermediate, parabasal, and basal zones.

In vaginal research, defining layers or regions accurately is paramount, as it precedes the characterization of their molecular and cellular attributes. Our study pinpointed several molecules as promising markers for distinguishing various zones. Notably, KRT13 and KRT5 were found to specifically label nearly all fetal vaginal epithelial cells, establishing them as epithelial markers (Figure 4C and 5B). Additionally, we identified distinct markers specific to different epithelial regions: IGSF9, NGFR, COL17A1, TP63, KRT14, and TP73 consistently labeled cells in the basal region; PI3 and TGM1 specifically targeted parabasal cells; CDH1 marked cells in both the basal and parabasal regions; SPINK7 and C15orf48 effectively identified cells in the intermediate region; while LCE3E, FLG, and KRT78 reliably labeled cells in both the intermediate and superficial regions.

Keratin subtypes, which are essential for both structural support and metabolic processes regulation, exhibit selective expression patterns in the human vagina. This heterogeneity is especially pronounced across cell types and spatial zones. Our comprehensive analysis of keratin expression patterns underscores the diversity within vaginal epithelial cells. Notably, KRT5/14 is most abundant in the basal zone, gradually decreasing towards the parabasal and intermediate zones (Figure 4C, 5A and 5C). This expression pattern parallels the regulation of epidermal stem cell proliferation, necessitating a relatively high level of KRT5/14 expression. In epidermal cells, the downregulation of K5/14 occurs as cells progress from the basal to suprabasal layers, a pattern observed in skin development [58]. This suggests that the human vaginal epithelium maintains stem cell proliferation in the basal zones while undergoing differentiation towards the superficial zone, mirroring a developmental pattern observed in the mouse vagina [12].

The vaginal epithelium, a squamous epithelium, relies on a renewal process to maintain its integrity and homeostasis. This renewal is hypothesized to involve the differentiation and proliferation of stem cells located in the basal region. Notably, the vaginal epithelium has demonstrated the capacity to fully recover from atrophy, a condition often characterized by the absence of mature differentiated keratinocytes in the upper layers of the vaginal epithelium [11, 59, 60], in response to estrogen [12]. This regenerative ability suggests the presence of a population of epithelial adult stem cells within the vagina. However, despite a few studies on vaginal stem cells, no clear epithelia stem cell marker has been identified for the human vagina [12]. In our data, the basal zone exhibited significant enrichment of pathways associated with cell cycle regulation, apoptosis, oxidative phosphorylation, the tricarboxylic acid (TCA) cycle, and various metabolic processes (Figure 6). This activity suggests a potential role for this zone in the maintenance, proliferation, and differentiation of human vaginal epithelial stem cells. Transcription factor TP63 is a marker for cells in the basal region of the vaginal epithelium. Rodent studies have shown that Tp63-expressing vaginal epithelial cells have latent skin competence, explaining the occasional observation of aberrant hair follicles or sebaceous glands in the vagina [40]. In the squamo-columnar region of the anorectal junction, a population of KRT17-positive basal cells demonstrates the capacity to sustain a squamous epithelium during normal homeostasis and actively contributes to the repair of a glandular epithelium following tissue injury [61]. Similarly, in the squamo-columnar cervical junction, KRT17-high P63+KRT5+ cervical cells show the ability to differentiate into both squamous and columnar cells [62]. In our vagina samples, KRT17 was uniquely observed in a subset of cells in the basal layer (Figure 5E), reflecting the heterogeneity of cells in this layer. Whether KRT17 is a marker for vaginal epithelial stem cells remains to be experimentally verified. The NGFR+AXIN2+ marker, identified as a stem cell marker in mouse vaginal epithelia [12], labels the fetal basal layer of the human vaginal epithelium (Figure 7A). Additionally, basal region markers such as IGSF9, COL17A1, KRT14, and TP73, along with the stem cell marker SOX2, are potential markers for human vaginal epithelial stem cells. Further research is needed to confirm the identity and function of these markers in the human vagina.

The immune function of vaginal epithelial cells is underscored by the prevalence of diseases related to microbial infections, emphasizing their crucial role as a barrier against pathogens. A particularly noteworthy finding is the spatially specific distribution of immune cells within the vagina (Figure 4D, S4B and S4C), shedding light on their contribution to maintain homeostasis across its various regions. Importantly, the basal zone also showed an enrichment in human immunodeficiency virus 1 infection and Human T-cell leukemia virus 1 infection (Figure 6), highlighting the immune function of vaginal epithelial cells. The parabasal zone demonstrated an elevated level of substance synthesis and metabolic activity, with a notable focus on lipids. This included Steroid biosynthesis, Linoleic acid metabolism, alpha-Linoleic acid metabolism, Ether lipid metabolism (Figure 6). The vaginal epithelium is characterized by a high lipid composition, with phosphatidylethanolamine, cholesterol, and phosphatidylcholine reported as dominant components [61]. These lipids are believed to contribute to the epithelium’s defense against pathogenic microorganisms and regulate its permeability to water and various chemical substances. Our analysis specifically points to an enrichment in Mucin type O-glycan biosynthesis within the intermediate zone, their biological role is unclear, may function for barrier.

Our focus on vaginal samples from 22-week-old fetuses has revealed striking molecular and histological parallels with reproductive-aged women, showcasing a stratified squamous epithelium high in glycogen content [62]. However, two sequencing samples from two 22^+^ week fetuses are limited, susceptibility to outcomes influenced by individuals. To paint a more nuanced and comprehensive picture, it is imperative to extend our investigations to a wider age spectrum and more samples. This approach will yield richer temporal and spatial insights, thereby refining our grasp of the vaginal epithelium’s intricate functions and developmental trajectories. Additionally, functional validations and targeted experiments are warranted to corroborate the inferences drawn from the spatial transcriptomics data.

In conclusion, our study elucidates the molecular landscape and heterogeneity of the human fetal vaginal epithelium. The integration of spatial transcriptomics, keratin profiling, and functional enrichment analyses provides a multi-faceted perspective on the cellular composition and activities within distinct epithelial zones. These findings not only advance our understanding of prenatal vaginal development but also carry significant implications for understanding adult epithelial characteristics and functions. This data also informs our understanding of squamous epithelial biology. Further explorations in this area holds the potential to unveil additional intricacies of vaginal tissue biology and its profound connections to health and disease.

## Acknowledgments

We would like to thank Ms. Ling Chen from the Pathology Department of Nanjing Drum Tower Hospital for her advice on fetal vaginal histology. We are grateful for the sequencing platform and bioinformation analysis of Gene Denovo Biotechnology Co., Ltd (Guangzhou, China). Thanks to Mr. Xiaodan Cai of Gene Denovo Biotechnology Company for his advice on spatial transcriptome data analysis.

## Fundings

This study was supported by research grants from the Strategic Priority Research Program of the Chinese Academy of Sciences (XDA16040101), National Key R&D Program of China (2021YFC2701603), National Natural Science Foundation of China (82271653) and Jiangsu Provincial Obstetrics and Gynecology Innovation Center (CXZX202229).

## Conflict of interest

The authors have declared that no conflict of interest exists.

## Author contributions

GFZ and YLH designed the experiment and collected the fetal vagina and performed the preprocessing for spatial sequencing. ZYY analyzed the 10x Visium data, performed the immunohistochemistry, Masson’s trichrome staining, immunofluorescence and wrote the manuscript. GFZ revised the manuscript. PPJ and ZRP collected the fetus for our study. QZ helped with bioinformatics analysis.

## Availability of Data and Materials

All available original data has been deposited. The raw sequence data presented in this study has been deposited in the Genome Sequence Archive (Genomics, Proteomics & Bioinformatics, 2021) hosted by the National Genomics Data Center (Nucleic Acids Res, 2022) at the China National Center for Bioinformation, affiliated with the Beijing Institute of Genomics, Chinese Academy of Sciences. The dataset, assigned the identifier GSA: HRA006835, is temporarily confidential but accessible. Furthermore, the Spaceranger output and HE staining maps corresponding to the sequenced samples documented in this paper have been archived in OMIX, also hosted by the China National Center for Bioinformation / Beijing Institute of Genomics, Chinese Academy of Sciences. The full dataset is accessible under the accession number OMIX005958 at https://ngdc.cncb.ac.cn/omix.

## Supplementary information for

**Figure S1.**
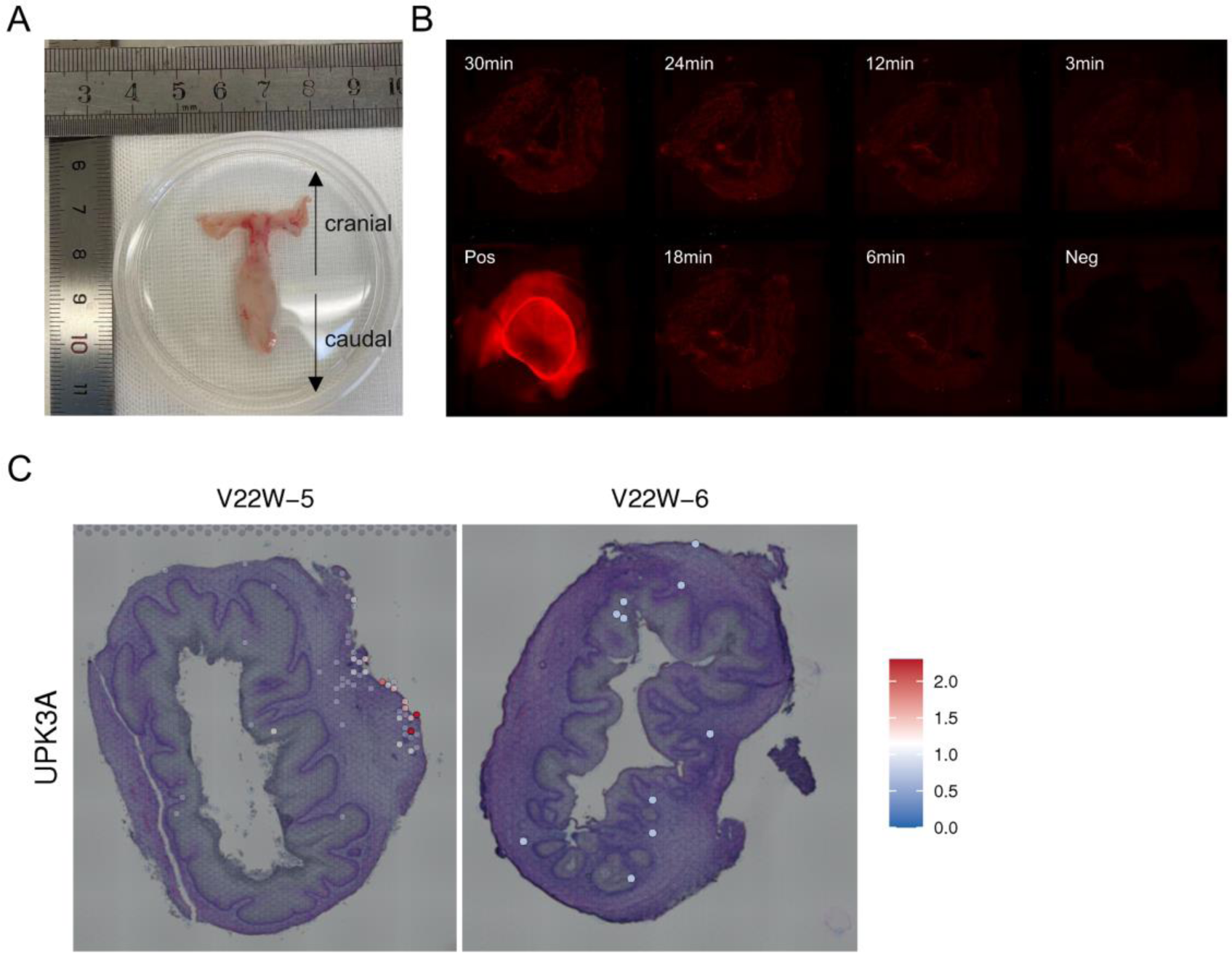
Spatial Transcriptomics Tissue Optimization. (A) Whole-mount photographs of a human female reproductive tract at 22+ weeks. (B) Optimization of permeabilization time, with positive (Pos) and negative (Neg) controls. (C) Spatial mapping of UPK3A indicating the region corresponding to the urethra, facilitating the removal of spots around this region to minimize interference from urethral genes.

**Figure S2.**
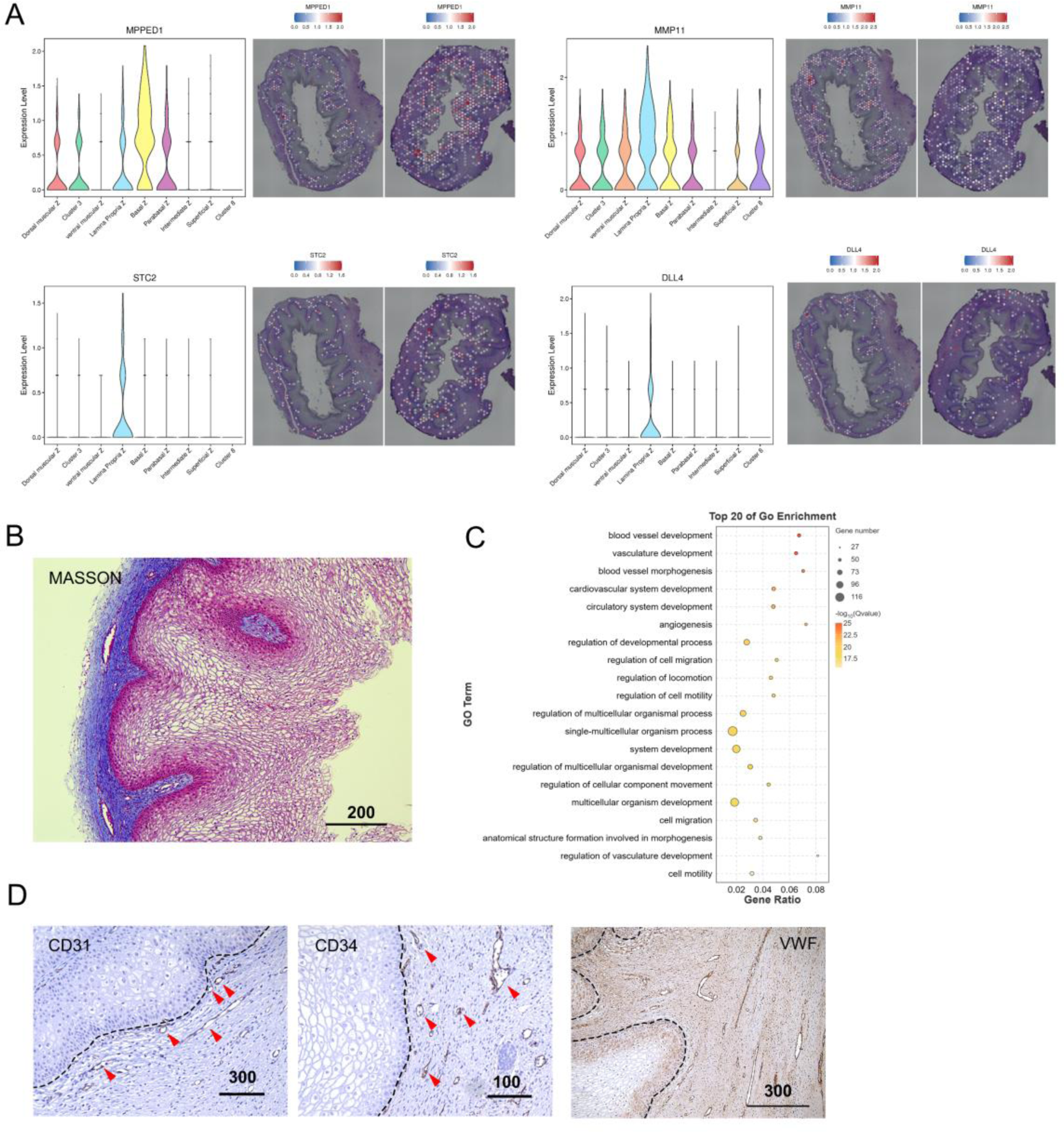
Cluster-Specific Genes Expressed in the Lamina Propria. (A) Violin plot (left) accompanied by spatial mapping (right) of DEGs in the lamina propria, showcasing the spatial distribution of specific molecules. (B) Masson staining of the fetal vaginal wall provides visualization of tissue composition. (C) Bubble plot displaying the top 20 Gene Ontology (GO) enrichments for the lamina propria. Threshold: adjust q-value < 0.05. (D) Immunohistochemistry (IHC) staining of blood vessel markers CD31, CD34, and VWF in the human fetal vagina, highlighting the vascular structures in the lamina propria.

**Figure S3.**
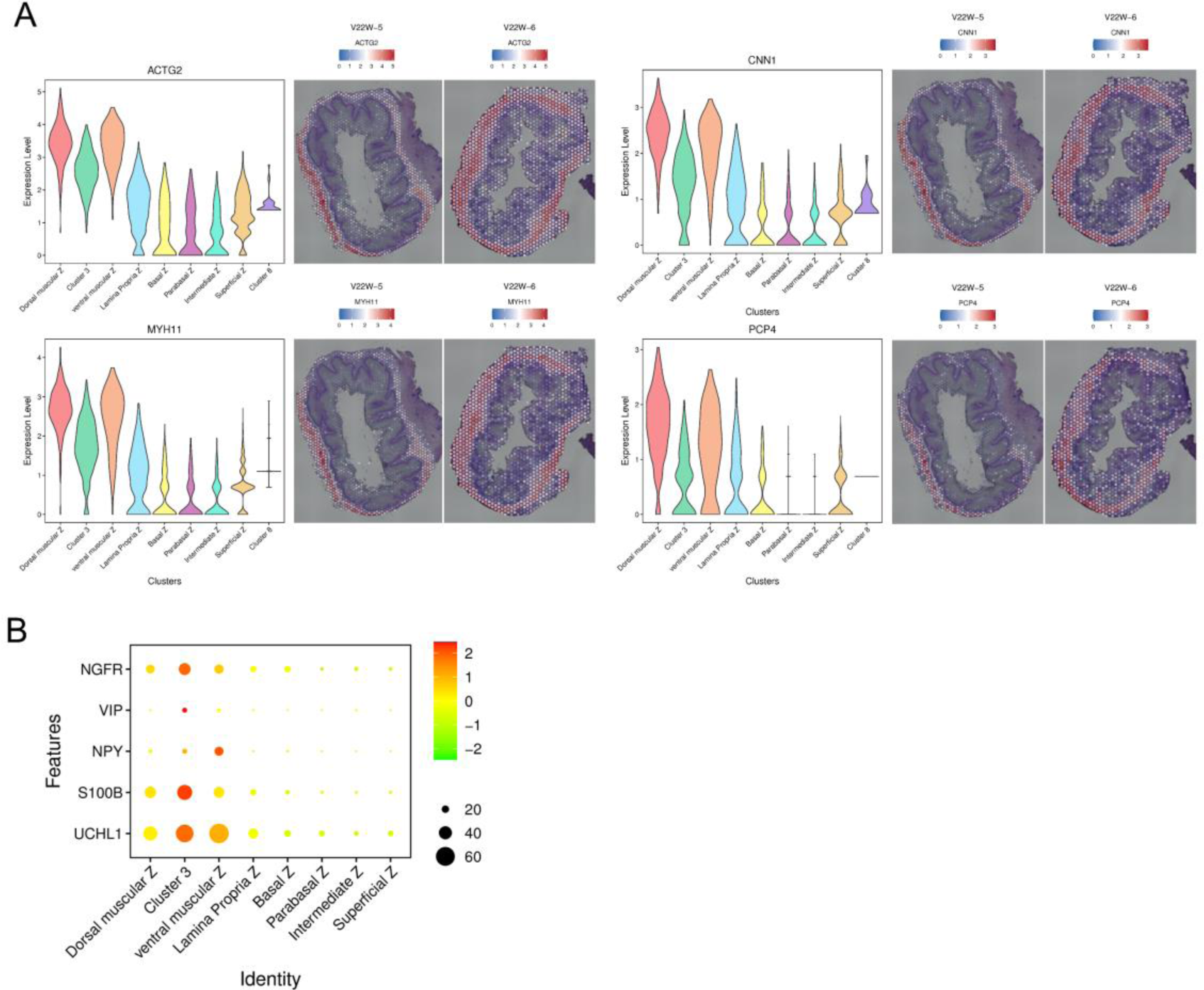
Cluster-Specific Genes Expressed in the Muscular Propria. (A) Left: Violin plot depicting differentially expressed genes (DEGs) in the muscularis propria. Right: Spatial mapping of these DEGs, illustrating their distinctive spatial distribution within the muscular propria. (B) Plot of neuron-related genes mapped against different vaginal regions, providing insights into their expression patterns in the context of the muscular propria.

**Figure S4.**
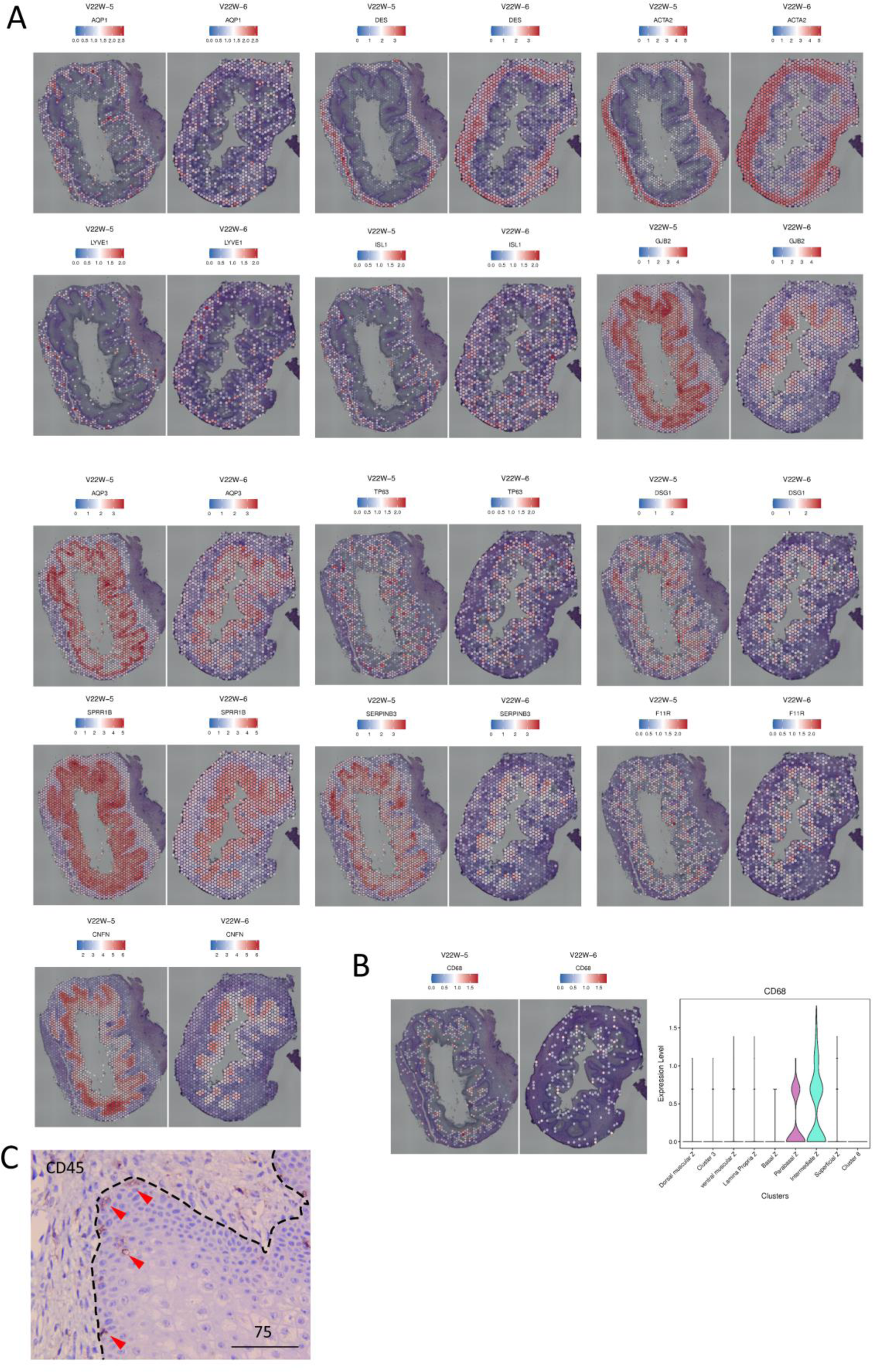
Tissue Mapping of Canonical Markers. (A) Spatial mapping of canonical markers in the H&E staining slices, providing an overview of their distribution within the tissue. (B) Dominant expression of CD68 in the parabasal and intermediate zones, as revealed by spatial mapping. (C) Immunohistochemical (IHC) staining depicting the presence of CD45+ cells in the basal zone of epithelia.

**Figure S5.**
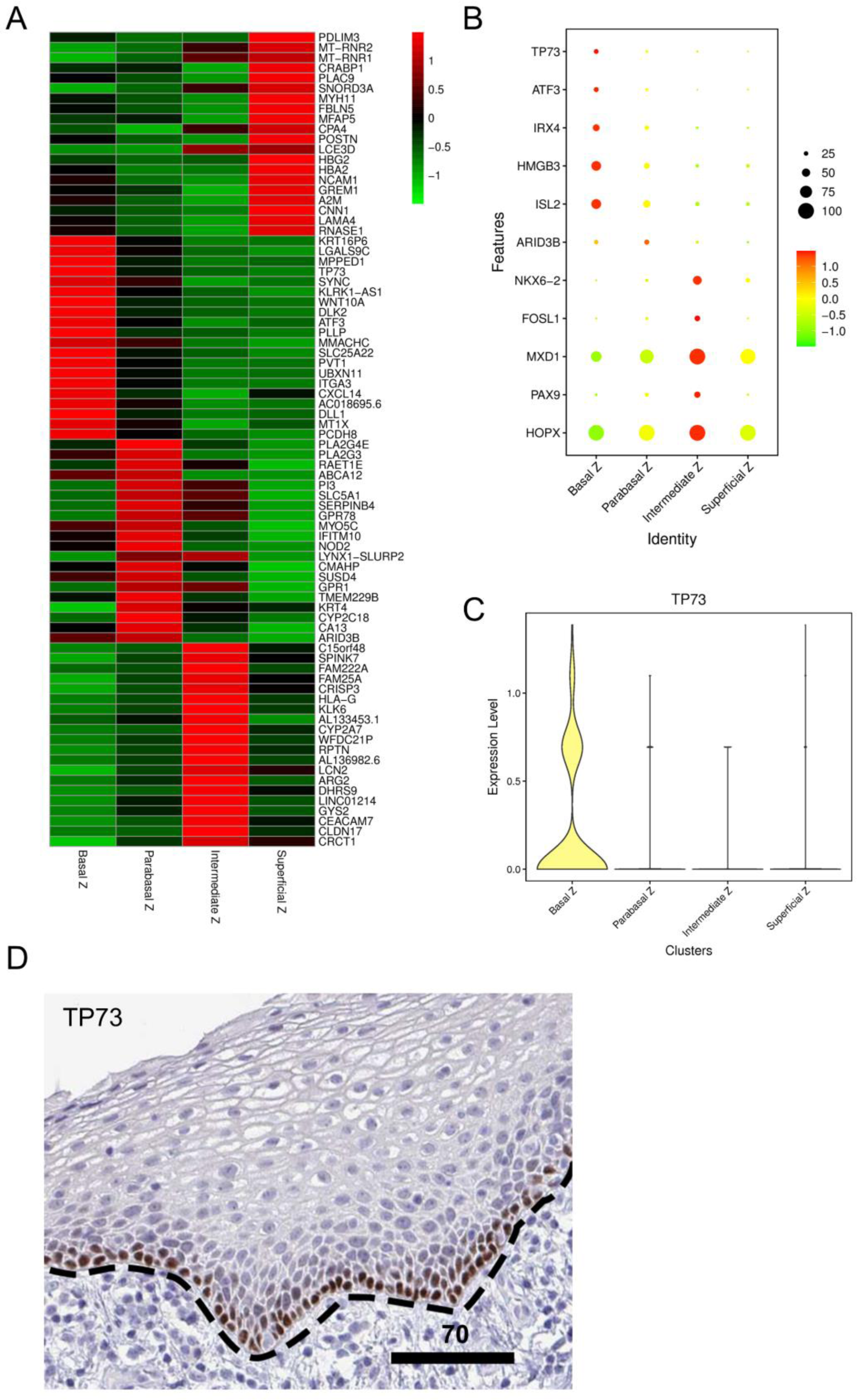
Identification of Differentially Expressed Genes in Vaginal Epithelium. (A) Heatmap illustrating the Differentially Expressed Genes (DEGs) among different zones of the vaginal epithelium. For clusters with fewer than 10 genes, all DEGs are displayed. (B) Bubble plot illustrating the top 5 transcription factors (TFs) associated with DEGs in different zones of the vaginal epithelium. All TFs are displayed for clusters with fewer than 5 TFs. (C) Violin plots displaying zone-specific transcription factor TP73 expression, focusing on the basal zone. (D) Immunohistochemical (IHC) staining of TP73 in adult vagina from the Human Protein Atlas (see Methods for details), revealing exclusive expression in the basal layer of the vaginal epithelium.

**Table S1.** Antibodies used for immunohistochemistry (IHC) and immunofluorescence (IF)

| Primary antibodies |  |  |  |  |
| --- | --- | --- | --- | --- |
| Antigen | Supplier | Cat. # | IHC | IF |
| ESR1 | abcam | ab23063 | 1:200 | NA |
| PGR | CST | #8757 | 1:200 | NA |
| CD31 | abcam | ab182981 | 1:100 | NA |
| CD34 | abcam | ab81289 | 1:200 | NA |
| KRT5 | abcam | ab52635 | NA | 1:200 |
| FLG | abcam | ab221155 | 1:1000 | NA |
| CDH1 | abcam | ab76055 | 1:400 | NA |
| TGM1 | proteintech | 12912-3-AP | 1:200-1:800 | NA |
| CD45 | abcam | ab10558 | 1:400 | NA |
| VIM | abcam | ab92547 | 1:200 | NA |
| KRT78 | abcam | ab122578 | 1:200 | NA |
| KRT17 | abcam | ab51056 | 1:200 | NA |
| MKI67 | abcam | ab279653 | NA | 1:1000 |
| MKI67 | abcam | ab15580 | 1:200 | NA |
| SOX2 | abcam | ab97959 | 1:500 | 1:500 |
| TP63 | Millipore | ABS552 | NA | 1:1000 |
| KRT5 | abcam | ab52635 | NA | 1:200 |
| VWF | abcam | ab6994 | 1: 200 | NA |
| Axin2 | abcam | ab107613 | 1:500 | NA |
| CK14 | abcam | ab119695 | 1:100 | NA |
| NGFR | abcam | ab52987 | 1:100 | NA |
| COL17A1 | ABclonal | A4808 | 1:100 | NA |
| INVOLUCRIN | ABclonal | A13311 | 1:100 | NA |
| CK13 | ABclonal | A0411 | 1:200 | NA |
| LCE3E | BT LAB | BT-AP03388 | 1:100 | NA |
| PGR | CST | #8757 | 1: 200 | NA |
| ESR1 | abcam | ab32063 | 1:200 | NA |

| Secondary antibodies |  |  |  |  |
| --- | --- | --- | --- | --- |
| Antibodies | Conjugation | Dilution | Supplier | Cat. # |
| Goat anti-rabbit | Rhodamine | 1/200 | Jackson ImmunoResearch | #111-025-045 |
| Goat anti-mouse | Rhodamine | 1/200 | Jackson ImmunoResearch | #115-025-003 |
| Goat anti-rabbit | Alexa Flur 488 | 1/200 | Jackson ImmunoResearch | #111-545-003 |
| Goat anti-mouse | Alexa Flur 488 | 1/200 | Jackson ImmunoResearch | #115-095-003 |

**Table S2.** Normalized Expression Levels of Keratin and Keratin-Associated Molecules in Human Fetal Vaginal Epithelium.

| GeneName | Basal Z | Parabasal Z | Intermediate Z | Superficial Z | Exp. Sum |
| --- | --- | --- | --- | --- | --- |
| KRT1 | 1.0452 | 0.6647 | 0.3219 | 0.4699 | 2.5017 |
| KRT2 | 0.0226 | 0.0146 | 0.0156 | 0.0015 | 0.0543 |
| KRT3 | 0.0198 | 0.0379 | 0.0344 | 0.0117 | 0.1038 |
| KRT4 | 44.9972 | 109.6414 | 63.0063 | 53.0220 | 270.6669 |
| KRT5 | 109.3333 | 56.1458 | 36.2563 | 66.9927 | 268.7280 |
| KRT6A | 211.2655 | 262.8659 | 179.9969 | 227.3994 | 881.5277 |
| KRT6B | 3.8616 | 5.9184 | 5.1594 | 5.7856 | 20.7249 |
| KRT6C | 41.2599 | 57.5102 | 41.4906 | 52.7783 | 193.0390 |
| KRT7 | 0.3729 | 0.2478 | 0.1906 | 0.2482 | 1.0595 |
| KRT8 | 3.1469 | 1.9184 | 1.3125 | 1.9280 | 8.3058 |
| KRT9 | 0.0847 | 0.0583 | 0.0563 | 0.0499 | 0.2492 |
| KRT10 | 2.8136 | 2.3061 | 1.5781 | 2.2643 | 8.9621 |
| KRT13 | 575.1017 | 705.4752 | 617.4750 | 621.2687 | 2519.3206 |
| KRT14 | 49.8757 | 18.0175 | 13.0813 | 25.0000 | 105.9744 |
| KRT15 | 5.7571 | 2.6501 | 1.7969 | 2.5932 | 12.7973 |
| KRT16 | 10.4605 | 10.9883 | 10.5188 | 11.1204 | 43.0880 |
| KRT17 | 2.5932 | 1.2799 | 2.2344 | 2.0793 | 8.1868 |
| KRT18 | 2.5960 | 3.0175 | 5.4313 | 3.1439 | 14.1887 |
| KRT19 | 30.0311 | 30.7930 | 37.0156 | 27.1248 | 124.9645 |
| KRT20 | 0.0000 | 0.0000 | 0.0031 | 0.0000 | 0.0031 |
| KRT23 | 0.0141 | 0.0175 | 0.0656 | 0.0191 | 0.1163 |
| KRT24 | 0.0141 | 0.0146 | 0.0156 | 0.0103 | 0.0546 |
| KRT25 | 0.0000 | 0.0058 | 0.0031 | 0.0000 | 0.0090 |
| KRT27 | 0.0028 | 0.0000 | 0.0000 | 0.0000 | 0.0028 |
| KRT31 | 0.0254 | 0.0262 | 0.0313 | 0.0294 | 0.1123 |
| KRT32 | 0.0085 | 0.0087 | 0.0063 | 0.0000 | 0.0235 |
| KRT33B | 0.0085 | 0.0204 | 0.0406 | 0.0088 | 0.0783 |
| KRT34 | 0.0056 | 0.0029 | 0.0094 | 0.0132 | 0.0312 |
| KRT36 | 0.0537 | 0.0437 | 0.0344 | 0.0176 | 0.1494 |
| KRT37 | 0.0000 | 0.0029 | 0.0031 | 0.0059 | 0.0119 |
| KRT40 | 0.0028 | 0.0029 | 0.0000 | 0.0000 | 0.0057 |
| KRT73 | 0.0028 | 0.0029 | 0.0031 | 0.0029 | 0.0118 |
| KRT74 | 0.0113 | 0.0175 | 0.0063 | 0.0044 | 0.0394 |
| KRT75 | 0.0169 | 0.0058 | 0.0063 | 0.0073 | 0.0364 |
| KRT76 | 0.0113 | 0.0000 | 0.0000 | 0.0044 | 0.0157 |
| KRT78 | 0.5960 | 1.1720 | 5.7688 | 2.9941 | 10.5309 |
| KRT79 | 0.0028 | 0.0029 | 0.0031 | 0.0015 | 0.0103 |
| KRT80 | 1.0424 | 1.8484 | 4.6063 | 2.6667 | 10.1637 |
| KRT81 | 0.0254 | 0.0058 | 0.0094 | 0.0117 | 0.0524 |
| KRT85 | 0.0000 | 0.0087 | 0.0000 | 0.0015 | 0.0102 |
| KRT86 | 0.0113 | 0.0000 | 0.0031 | 0.0044 | 0.0188 |
| KRT222 | 0.0056 | 0.0029 | 0.0000 | 0.0000 | 0.0086 |
| KRTAP11-1 | 0.0113 | 0.0350 | 0.0938 | 0.0264 | 0.1665 |
| KRTAP13-2 | 0.0169 | 0.0029 | 0.0031 | 0.0162 | 0.0391 |
| KRTAP1-5 | 0.0028 | 0.0000 | 0.0000 | 0.0015 | 0.0043 |
| KRTAP17-1 | 0.0113 | 0.0000 | 0.0094 | 0.0000 | 0.0207 |
| KRTAP19-1 | 0.0000 | 0.0000 | 0.0000 | 0.0029 | 0.0029 |
| KRTAP2-3 | 0.0028 | 0.0000 | 0.0000 | 0.0029 | 0.0058 |
| KRTAP3-1 | 0.0028 | 0.0029 | 0.0000 | 0.0000 | 0.0057 |
| KRTAP5-10 | 0.0169 | 0.0029 | 0.0125 | 0.0044 | 0.0368 |
| KRTAP5-7 | 0.0000 | 0.0029 | 0.0000 | 0.0000 | 0.0029 |
| KRTAP5-8 | 0.0028 | 0.0029 | 0.0000 | 0.0000 | 0.0057 |
| KRTAP5-9 | 0.0169 | 0.0087 | 0.0031 | 0.0088 | 0.0376 |

## Notes

https://ngdc.cncb.ac.cn/omix

https://ngdc.cncb.ac.cn/gsa/

https://www.proteinatlas.org/

### Competing Interest Statement

The authors have declared no competing interest.

